# Human Migration and the Spread of the Nematode Parasite *Wuchereria bancrofti*

**DOI:** 10.1101/421248

**Authors:** Scott T. Small, Frédéric Labbé, Yaya I. Coulibaly, Thomas B. Nutman, Christopher L. King, David Serre, Peter A. Zimmerman

## Abstract

The human disease lymphatic filariasis causes the debilitating effects of elephantiasis and hydrocele. Lymphatic filariasis currently affects the lives of 90 million people in 52 countries. There are three nematodes that cause lymphatic filariasis, *Brugia malayi, B. timori*, and *Wuchereria bancrofti*, but 90% of all cases of lymphatic filariasis are caused solely by *W. bancrofti*. Here we use population genomics to identify the geographic origin of *W. bancrofti* and reconstruct its spread. Previous genomic sequencing efforts have suffered from difficulties in obtaining Wb DNA. We used selective whole genome amplification to enrich *W. bancrofti* DNA from infected blood samples and were able to analyze 47 whole genomes of *W. bancrofti* from endemic locations in Haiti, Mali, Kenya, and Papua New Guinea. Our results are consistent with a Southeast Asia or East Asia origin for *W. bancrofti* spread around the globe by infecting migrating populations of humans. Austronesians probably introduced *W. bancrofti* to Madagascar where later migrations moved it to continental Africa. From Africa, *W. bancrofti* spread to the New World during the transatlantic slave trade. The greater genetic diversity of *W. bancrofti* populations from Haiti are also consistent with genetic admixture from multiple source populations. Genome scans for locally adapted haplotypes identified genes associated with human immune suppression and insecticide sensitivity. Locally adapted haplotypes may provide a foundation to understand the distribution of *W. bancrofti* compared to that of other filarial nematodes and how populations may differ in response to eradication efforts.

## Introduction

Infectious diseases have shaped the history of the human populations and are still today one of the main burdens on human health. In addition to clinical studies and laboratory investigations, knowledge of the evolutionary history of a disease can improve our understanding of the current threat of infectious diseases, the immune response of the host to the pathogen or the infection dynamics. The evolutionary history of a pathogen may also provide clues to virulence or vector adaptation. In the event of a population bottleneck or expansion of the disease, identifying the driving factors may bring insight into control or future risk.

Lymphatic filariasis is a human disease currently infecting over 90 million people across 52 countries and is the second leading cause of permanent and long-term disability worldwide [Ottesen et al., 2008, World Health Organization, 2010, World Health Organization, 2012]. Disability is associated with recurrent adenolymphangitis, hydrocele, and elephantiasis that result in a loss of 5.9 million disability-adjusted-life-years [Gyapong et al., 2005, World Health Organization, 2015, World Health Organization, 2016]. Current treatment involves the drug Diethylcarbamazine (DEC), although DEC can cause serious complications (including encephalopathy and death) in patients who may also have onchocerciasis (caused by infection with *Onchocerca volvulus*) or loiasis (caused by *Loa loa*). Alternative use drugs, such as ivermectin, kill only the larval blood-stage (microfilariae), but not the adult worms thus have limited long-term effects on the worm populations [Centers for Disease Control and Prevention, 2018].

*Wuchereria bancrofti* (Wb) is responsible for over 90% of the cases of lymphatic filariasis (World Health Organization 2015) and is widely distributed throughout the tropics. The closely related nematodes *Brugia malayi* and *B. timori* are responsible for the remaining cases of lymphatic filariasis. *B. malayi* is found throughout Southeast Asia and southern India, while *B. timori* is found only in parts of Indonesia [World Health Organization, 2015]

Wb is highly specialized for human hosts and efforts to culture Wb *in vivo* in silver-leaf monkeys (*Trachypithecus cristatus*; [Palmieri et al., 1983]) or *in vitro* have failed to produce viable adult worms [Zaraspe and Cross, 1986, Franke et al., 1987, Franke et al., 1990]. Much of what we know about Wb biology is derived from studies of *B. malayi* in rodent hosts. However, the vast difference in distribution, host preference, and disease incidence between filarial nematodes demonstrate a need to investigate the evolutionary history of Wb specifically in an effort to understand why it is so widespread in relation to other filarial nematodes.

In light of experimental limitations, population genomics could offer an alternative approach to study Wb but is hampered by insufficient DNA quantity and the presence of multiple parasite genomes present in a single infected individual. In a previous study of Wb, we used individual Wb worms dissected from mosquitoes but the low infectivity rate of Wb in mosquitoes (2%; [Paily et al., 2009]) means that most captured mosquitoes will not contain Wb limiting the number of samples available. However, using whole genome amplification to create sufficient template for sequencing, we showed nucleotide diversity in the sampled population of Papua New Guinea was 0.001 [Small et al., 2016] similar to diversity in populations of *O. volvulus* [Choi et al., 2016]. However, little else about the history of Wb could be inferred from only the single population.

Here we use selective whole genome amplification (sWGA) to specifically enrich Wb DNA while avoiding amplification of Human DNA present in a human blood sample [Leichty and Brisson, 2014, Clarke et al., 2017]. sWGA allows us to enrich DNA from a single microfilaria (the transmitted stage of the parasite found in human blood) isolated from preserved blood samples collected prior to mass drug administration. We used sWGA to generate 42 new genome sequences from Wb worms originating from multiple endemic areas in Africa, Papua New Guinea, and Haiti and reconstruct the evolutionary and demographic history of Wb. Our specific goals were to 1) identify the geographic origin of Wb and thus lymphatic filariasis, 2) reconstruct how Wb spread from its geographic origin to other parts of the world, and 3) identify population-specific adaptations that may provide fitness advantages for Wb in the different endemic environments.

## Results

### Improved *W. bancrofti* genome sequence and gene prediction

The Wb genome sequence (PRJNA275548) was assembled from short reads and contains 5,670 contigs and 5,105 scaffolds [Small et al., 2016]. We used long reads generated with PacBio sequencing to increase the assembly and infer potential chromosome homology with other filarial nematodes. PacBio sequencing of a WGA library prepared from a dissected L3 Wb worm (see methods) produced 1.2 million sub-reads with an average of 5.0 kb in length. PacBio reads were used only for scaffolding the previous assembly and resulted in 856 scaffolds, with an N50 of 537 kb and with 198,000 introduced Ns. Further scaffolding using synteny with ragout v1.7 resulted in 17 scaffolds longer than 100 kb with an N50 of 12.37 Mb and with 1.4 Mb of introduced Ns. The final assembly total size was 88.46 Mb with the longest scaffold of 24.2 Mb (supplementary table S1). Visualization of the scaffolding and comparisons to *B. malayi* for the 17 largest contigs (length > 100 kb) are presented in supplementary figure S1)

To aid in gene predictions, we also sequenced an RNA library generated from filtered microfilaria preserved in RNALater (see methods in [Erickson et al., 2013]). Sequencing generated 600 million reads of which 1.7% mapped to the Wb genome and 98.3% mapped to Hg19. Using these data, we predicted 9,651 genes, including 9,517 assigned to the 17 longest contigs.

### Genome sequencing for 42 individual worms

One limitation to genomics in Wb is the minute amount of DNA contained in a single individual worm’s genome. We used the program sWGA [Clarke et al., 2017] and amplified ten microfilariae from each of 26 infected human blood samples from Mali (N=5), Papua New Guinea (PNG) (N=11), Kenya (N=5), and Haiti (N=5) (Supplementary Materials). After amplification, we assessed the ratio of Wb to human DNA and selected 42 individual worms with the best ratio for sequencing on 12 lanes of Illumina HiSeq X. After quality control and adapter trimming, we retained, on average, 40 million reads per individual worm with 90% of reads mapped to the improved Wb genome.

We compared these 42 newly sequenced genomes with 13 previously sequenced genomes from Papua New Guinea [Small et al., 2016] and a single genome from Mali [Desjardins et al., 2013]. We removed individuals that were full-siblings as individuals identified in Small et al. 2016 as well as individuals with high heterozygosity across the mitochondrial genome (suggesting that the sequences derive from multiple individuals). This led to a total of 56 worms for structure analysis, with 47 worms selected to limit missing data for downstream analysis (supplementary table S2). Final samples sizes were: Haiti=7, Mali=11, Kenya=9, PNG=20.

We identified 403,487 SNPs among the 47 Wb individuals, with an average Ts/Tv ratio of 3.58 and average coverage per called genotype of 141 X (all positions with < 10 X were masked in our analyses). 86% of the SNPs were found in intergenic regions of the genome (enrichment of 1.16). 9% of SNPs were found in introns (enrichment of 0.53). The remaining 5% of SNPs were found in predicted protein-coding regions (enrichment of 0.60) (supplementary table S3).

### Genetic Diversity is similar among *W. bancrofti* populations

The population closer to the geographic origin of Wb are likely to harbor greater genetic diversity than founder populations (due to successive bottlenecks in the history of the latter). The median value of genetic diversity was not statistically different among all populations (Haiti=0.00084, Mali=0.00072, Kenya=0.00076, and PNG=0.00074, supplementary figure S2A). While the population of Haiti had the highest variance in genetic diversity, this may be due to multiple admixture events (see below). However, the recent estimated effective population size (Fig 2B., table 1) is largest in Papua New Guinea (PNG), and it is likely that the geographic origin is in Southeast Asia.

**Table 1:** Demographic Parameter Estimates.

| Population | Effective size | Divergence (yrs) |
| --- | --- | --- |
| Haiti | 50-1,100 | – |
| Mali | 30-1,200 | – |
| Haiti-Mali | 8,318-32,000 | 394-502 |
| Kenya | 100-6,000 | – |
| Haiti-Mali-Kenya | 697-2,500 | 1,450 – 1,898 |
| Papua New Guinea | 500-10,000 | – |
| Haiti-Mali-Kenya-PNG | 98,000-125,000 | 37,000-57,000 |

**Figure 2:**
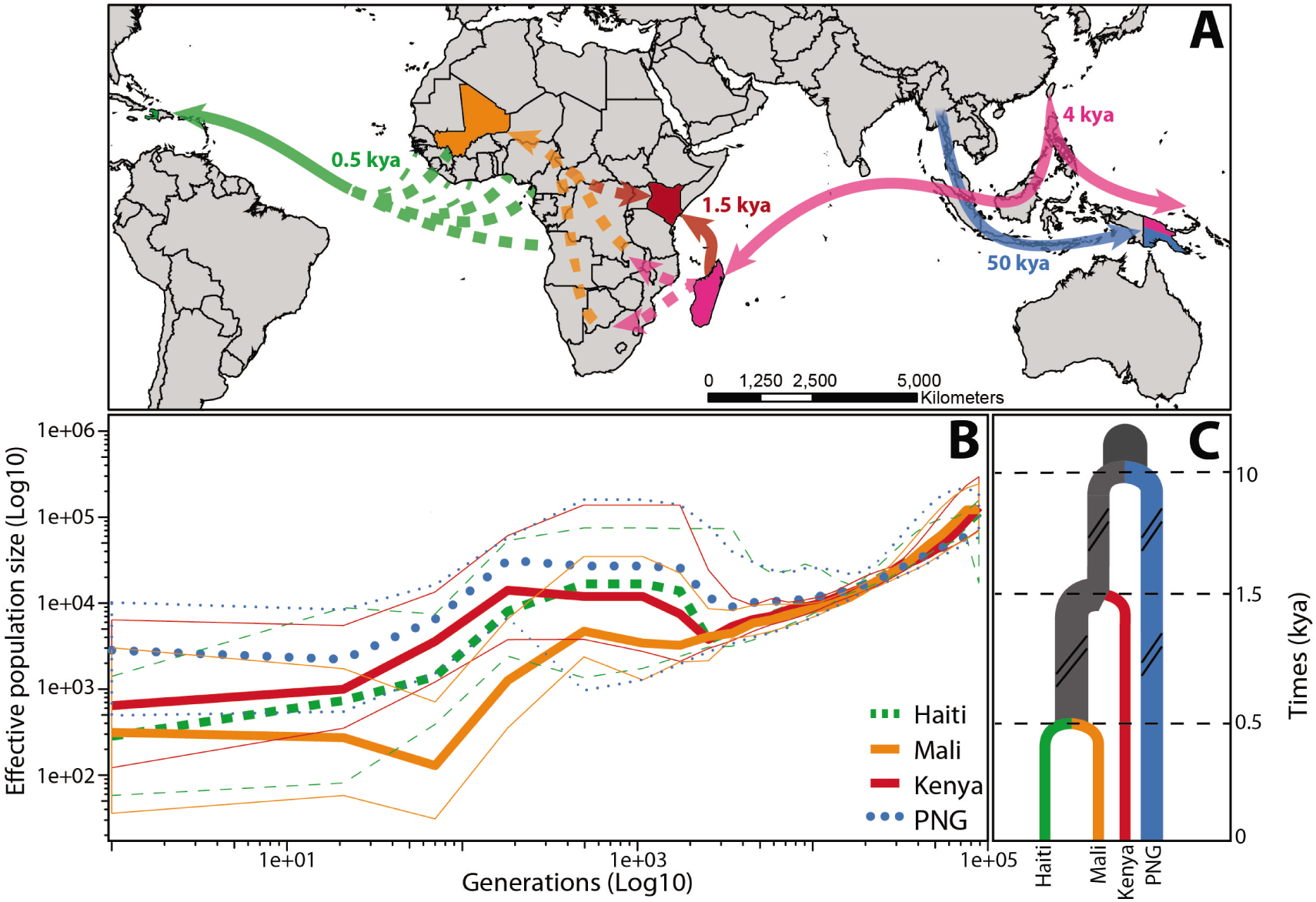
Demographic history of *W. bancrofti*. A) Map representing potential routes of Wb dispersal highlighting sampled populations and shared ancestry: Haiti (green), Mali (orange), Kenya (red), Papua New Guinea (blue). Pink colors represent unsampled and thus inferred population ancestry connecting Wb populations (color figure available online). B) Reconstruction of change in effective population size for each Wb population using MSMC2 and PopSizeABC. Colors denote population and thinner shaped lines of same shape represent 95% confidence intervals for the effective population sizes. C) Cartoon representing the highest likelihood model for Wb demographic history. Times along the vertical axis represent thousands of years before present (ago).

### Population bottlenecks in African and Haitian *W. bancrofti* populations

The Tajima’s D value summarized across the whole genome reflect the demographic history of one population (i.e., negative for rapidly expanding populations and positive for populations undergoing a contraction). The median value of Tajima’s D was similar among African and Haitian populations (Mali=0.4202, Kenya=0.2446, Haiti=0.3159) with a median value greater than 0. By contrast, the median value of Tajima’s D in PNG was -0.2151; supplementary figure S2B).

The scaled site frequency spectrum agreed with the median Tajima’s D values, with an excess of intermediate variants in the Haitian and African populations. The trajectory is relatively flat for PNG; an indication of stable and constant population size ([Miles et al., 2017]; supplementary figure S2C).

The correlation among genotypes followed this same expectation, with higher linkage disequilibrium in the African and Haitian populations compared to the PNG population (supplementary figure S2D). These trends indicate that African and Haitian populations have likely undergone recent population contractions, founder effects, while the PNG population has likely remained constant.

### Population differentiation is greatest between Papua New Guinea and African populations

Population differentiation should be smaller between recently diverged populations or populations exchanging migrants. Principal component analysis separated individuals according to their geographical origins (fig. 1A; PCA for each chromosome in supplementary figure S4 & S5). Interestingly, PC1, that explained 7.5% of the variance, separated PNG Wb from all other MF. Phylogenetic reconstruction, using genome-wide SNPs, confirmed these results and assigned worms into reciprocally monophyletic groups in regards to geographic location (fig. 1C). PCA also indicates that the PNG sample is genetically distinct from other regions, while the African and Haitian populations are similar due to being more recently diverged or ongoing gene flow.

**Figure 1:**
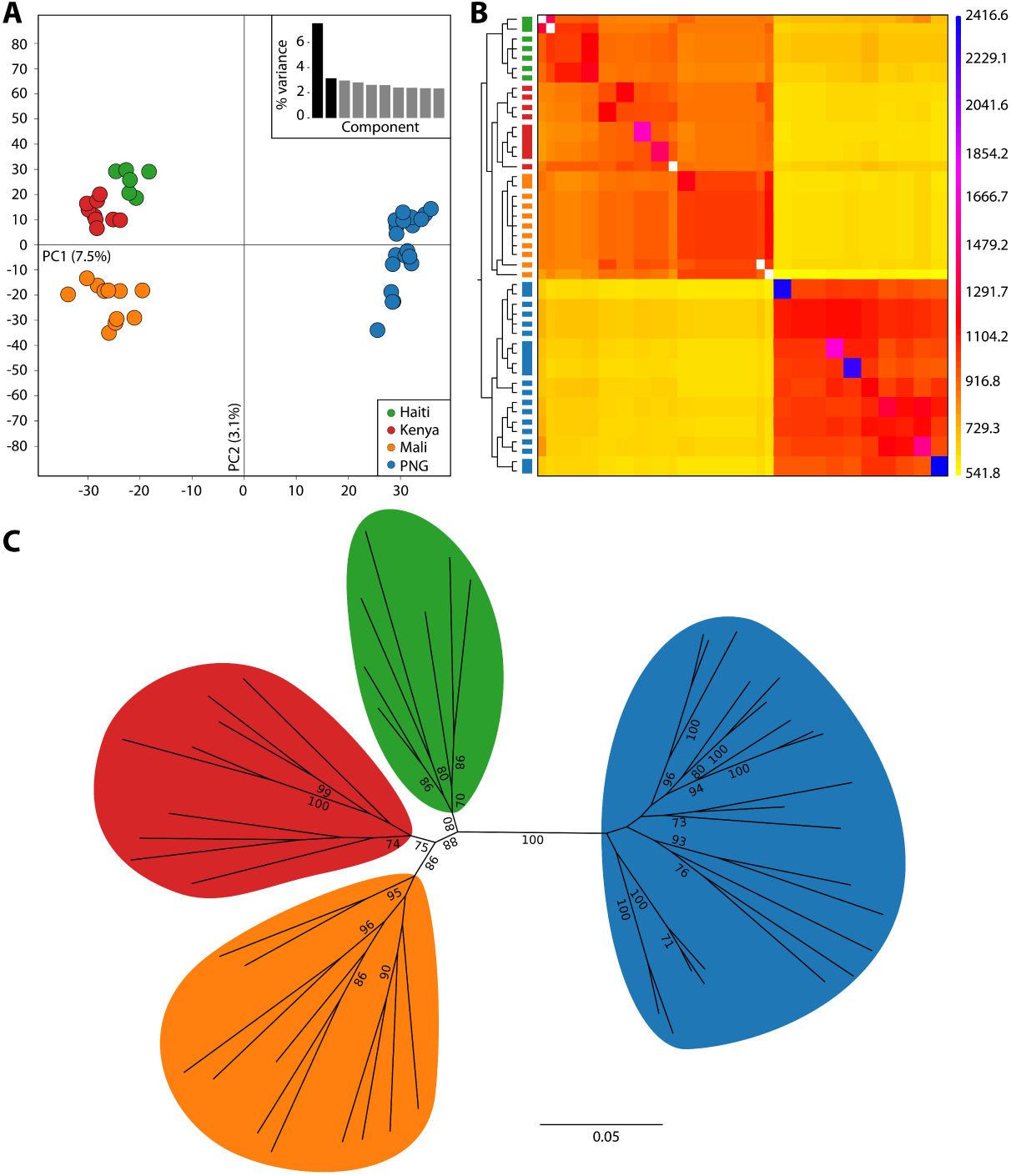
Genetic structure and ancestry among *W. bancrofti* populations. A) Principal Components Analysis (PCA) of 10,000 SNPs across the genome after removing sites in high LD. PCA 1 separates Haiti (green), Mali (orange), Kenya (red) from the population of Papua New Guinea (blue). PCA 2 separates Kenya, Mali, and Haiti. Percent variation explained by each component is summarized in the bar plot (inset). B) Coancestry matrix (a summary of nearest neighbor haplotype relationships in the data set) of Wb populations using fineSTRUCTURE. Trees relating individuals to the co-ancestry matrix are along the vertical axis with populations denoted by colors corresponding to PCA. Fused boxes of similar color denote worms sampled from the same host infection with high coancestry. A plot with individual labels on the vertical axis is available in supplementary figure S3. C) Whole genome SNP phylogeny using SNPhylo highlights that individuals from the same population are monophyletic. Support is shown for bootstrap values greater than 70.

### Partial admixture between African and Haitian populations of *W. bancrofti*

Using ADMIXTURE to allow for partial assignment to a number of ancestral populations, we found the highest support for two ancestral clusters, one with all worms from PNG and a second with worms from Africa and Haiti (supplementary figure S6). We further tested individual ancestry by increasing the number of clusters, to determine if a hierarchical structure within each cluster may be a function of geography. This did not further sub-divide clusters but instead resulted in partially mixed ancestry between African and Haitian worms. Although these configurations had lower likelihoods than the initial clustering, they hint at more complex and partial ancestry between African and Haitian worms, consistent with the founding of the Haitian population from multiple sources (supplementary figure S7).

### Within infection genetic relatedness is highest in Mali

Parasites can be considered to have a hierarchical structure that consists of first all the parasites within an infected host (infra-population) and then the greater community across infected hosts. To determine if genetic relatedness was higher within host infections than among hosts within the same regions, we used chromosome level ancestry analysis in the program fineSTRUCTURE. We observed higher co-ancestry at the level of the host infections especially in the Mali population (linked, filled boxes fig. 1B). Higher relatedness may be due to inbreeding or local stratification. We also observed worms from the same infections clustering with worms from other infections, which may represent between host transmission (supplementary figure S3).

### Lack of support for post-divergence gene flow

Both gene flow and genetic drift influence the estimate of population divergence times. Since we are interested in estimating divergence times, we tested the fit of alternative models of divergence with and without post-divergence migration. We used *∂*a*∂*i to model the shared demographic history between each pair of Wb populations. The best fit model to the data was a strict isolation model without migration between populations after divergence (Likelihood Ratio Test of rejecting the null model of no migration, pvalue=0.30). Multiple population models evaluated in ABLE similarly supported a model with no post-divergence gene flow (AIC=36.20, supplementary table S4). Thus, we used a model with no post-divergence gene flow for the remainder of our analyses and when generating simulations for hypothesis testing (table 1).

### Most recent common ancestor supports a Southeast Asia origin of *W. bancrofti*

The oldest node, representing the most recent common ancestor of all populations, provides some insights on the geographic origin of Wb. Divergence times estimated from the strict isolation model support that the four Wb populations last shared a common ancestor 50 kya (thousand years ago; assuming one generation per year) with an ancestral effective population size of 98,000 -125,000.

### Divergence timing between *W. bancrofti* populations are consistent with known human migration times

Human migration proceeded out of Africa to colonize other parts of the world with the first humans reaching Southeast Asia around 40-50 kya [Wollstein et al., 2010, Nielsen et al., 2017]. The only known ancestral human migration from Southeast Asia back to Africa was the during the Austronesian migration [Bellwood et al., 2006]. The most recent common ancestor between our African Wb populations is 1.4 – 2.0 kya with an ancestral size of 697 – 987. The estimated time of the Austronesian arrival in Madagascar was 300 C.E. [Bellwood et al., 2006, Gray et al., 2009] (Fig 2). Given the similarity of these times and the lack of alternative hypotheses (see Discussion), Wb likely spread to Africa from Southeast Asia or East Asia during the Austronesian migration. The presence of Wb in the New World is more likely due to recent migrations since colonization along with the peopling of Americans would present an older divergence time (13 kya; [Skoglund and Reich, 2016, Nielsen et al., 2017]). The divergence time between the Mali and Haitian populations of Wb is 0.3 – 0.5 kya. The timing of the transatlantic slave trades, where Western and Central Africans were forcibly transported to the New World, was from early 1400 C.E. to late 1700 C.E. (estimates of 4,000 to Jamaica in 1518 C.E., [Wynter, 1984]). This is concordant with the divergence times between Mali and Haitian populations of Wb and it is likely that the Wb population of Haiti was founded by migrations from both Central and Western African populations of Wb (Fig 2, table 1).

### Serial founder effects reduced population sizes in African and Haitian populations

Under a model of serial founder events, a Wb population originating in Southeast Asia and traveling to Africa and then America, we would expect to observe bottlenecks consistent with a reduced population size. The historical trajectories of each Wb individual populations, estimated using both a coalescent approach [Malaspinas et al., 2016] and the genome-wide site frequency spectrum [Boitard et al., 2016] supported a continued decline in Wb population sizes from Africa and Haiti (Fig 2B). The Wb populations of Africa and Haiti underwent bottlenecks starting with the divergence from PNG (10 kya), after which their ancestral populations declined to an effective size of 2,000. The ancestral effective size of Haiti-Mali was 10 to 50 times larger than the ancestral size of Mali-Kenya, indicating a likely population expansion across Africa (Fig 2A, 2C).

The trajectories of Mali, Kenya, and Haiti diverge between 0.15-0.30 kya, with the Malian Wb population declining from an effective population size of 2,000 down to 300. Kenyan and Haitian Wb populations follow similar but not identical trajectories with an initial increase in size to 10,000 followed by a decrease to an effective size of 700 and 300, respectively (fig. 2B, table 1).

### Local adaptation in populations of *W. bancrofti*

As Wb worms spread to different regions, they would have come into contact with novel environments, including new mosquito vectors, co-transmitted pathogens, and different human immune responses. Uncovering locally adapted genes may provide clues as to why Wb has been successful at colonizing and infected human populations (e.g., in contrast to *B. malayi* and *B. timori*). We used haplotype-based analysis [Fariello et al., 2013] to identify regions of the genome where allele frequencies significantly differ between populations. We identified nine regions of the genome with signals of local adaptation (fig. 3). Functional annotations revealed that the genes in these regions are associated with human host interaction, reproduction, and chemical toxicity (table 2). Five genomic regions were consistent with adaptation in the Haiti Wb population, containing a total of five genes with few known functional annotations. A single gene was annotated, coding for cystatin, a proteinase inhibitor, related to host immunity [Schönemeyer et al., 2001, Zang and Maizels, 2001, Murray et al., 2005, Wang et al., 2017]. Seven regions were significant outliers in the Mali Wb population, containing five genes associated with host immune regulation (e.g. D-dopachrome tautomerase (DDT) as a co-factor of macrophage inhibitory factor (MIF-1) [Merk et al., 2011] and reproductive timing, nurf-1 [Large et al., 2016]. We also identified signals consistent with local adaptation at a GABA transporter previously noted as being upregulated in studies of the anthelmintic flubendazole [Jiang et al., 2005, Mullen et al., 2006, Casida and Durkin, 2013, O’Neill et al., 2016]. Only two regions, with a single known gene annotation, were consistent with adaptation in the Kenyan Wb population: the SLO-1 gene is a calcium-activated potassium channel [Welz et al., 2011]. Three other genes were possibly selected in both the Malian and Kenyan populations of Wb: Acetylcholinesterase (AChE), specifically noted as resistance to inhibitors of AChE (as identified in Caenorhabditis elegans; [Nguyen et al., 1995]), phosphatidate cytidylytransferase [Narayan et al., 1989, Crowther et al., 2010], and PAN domain [Casida and Durkin, 2013, Verma et al., 2017, Sundaraneedi et al., 2018]. While these findings will need to be validated experimentally, these genes largely associate with human-host interactions, worm reproduction, and chemical toxicity.

**Table 2:**
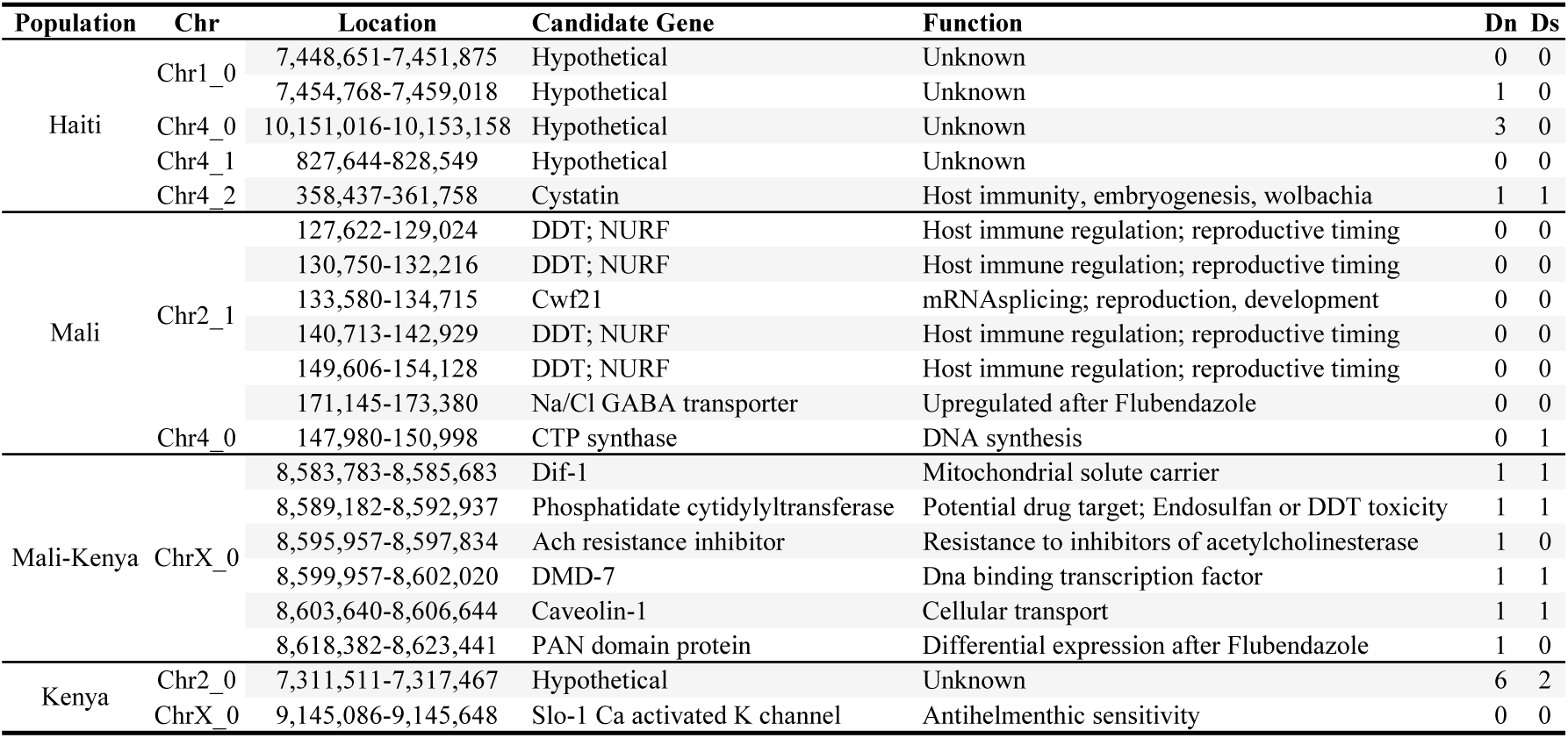
Results Selection Scan.

**Figure 3:**
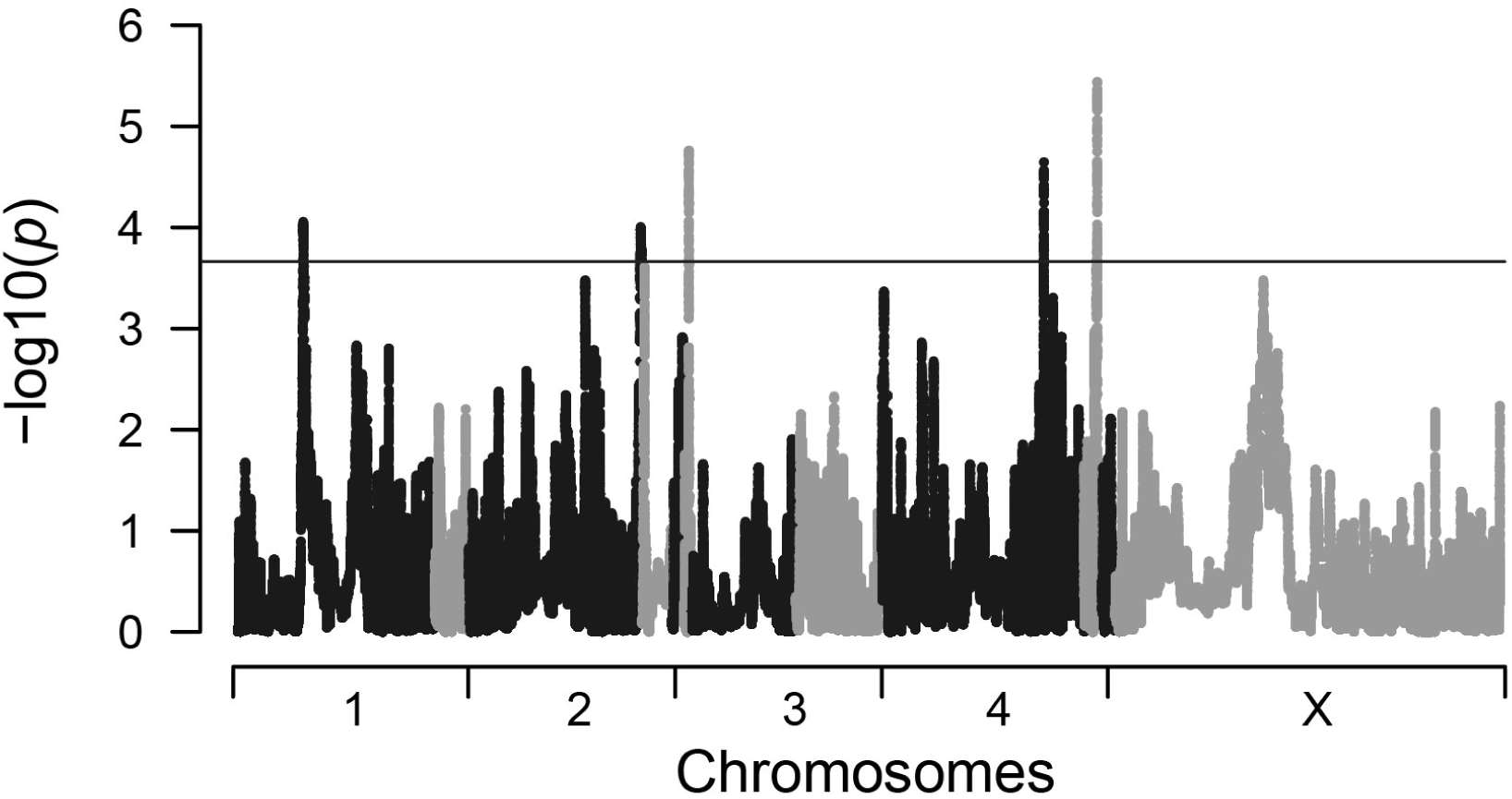
Manhattan plot of local adaptation in *W. bancrofti*. Results of selection scans using hapFLK to identify haplotypes that are present at high frequency in Wb populations. Analysis represents combined result of Haiti, Mali, and Kenya. Papua New Guinea was used as an outgroup for the analysis. Populations with signals of local adaptation were identified using eigen analysis. Details of specific genes are available in table 2.

## Discussion

The goals of our study were to infer the geographic origin of Wb (and thus of lymphatic filariasis), to determine how Wb spread from its origin to other regions, and identify any genes that were locally adapted within each endemic population. Using genomic data from 47 Wb worms, we inferred divergence times and effective population sizes consistent with a species origin in Southeast Asia. Based on our results, we hypothesize that Wb then spread from Southeast Asia to Africa by the Austronesian migration and later to the New World by the transatlantic slave trade. Our results taken as a whole increase our understanding of the evolutionary history of Wb and provided genomic resources for studying human – parasite evolution as well as evaluating ongoing elimination programs.

### Geographic Origins of *W. bancrofti*

We used population genetic models and whole genome summary statistics to test models of population divergence, migration, and admixture in an effort to infer the species origin of Wb (supplementary figure S8; table 1). The best fit model did not include post-divergence migration or recent migrations between pairs of populations (supplementary table S4). Our data revealed that the split between PNG and all other populations predates the divergence between African and Haitian populations of Wb, supporting a hypothesis of a Southeast Asian or East Asian origin.

Others have proposed a Southeast Asia origin for Wb as being consistent with both species diversity and archaeology [Laurence, 1989, Hoeppli, 1969, Fagg, 1977, Laurence, 1968]. The previous hypothesis speculated that the species origin of Wb is likely present-day Indonesia, because Indonesia is the endemic range of the three recognized Brugia species (*B. malayi, B. timori*, and *B. pahangi*) as well as the only other known Wuchereria species, *W. kalamantani* (parasite of silver-leaf monkeys; [Palmieri et al., 1980]). Even with genomic data, the origin of Wb is difficult to determine since we currently do not have samples from anywhere in the region except PNG. However, our current data is consistent with the hypothesis that the origin of Wb is in Southeast Asia or Indonesia. Further confirmation would require genomic samples of Wuchereria as well as Brugia from multiple populations in Indonesia and China. Although currently there are no publicly available population datasets for any wild Brugia species.

### How did *W. bancrofti* first arrive on New Guinea?

Our best fit demographic model included a single Wb colonization 50 kya followed by a later admixture 5 kya with the Austronesian migration. We hypothesize that the original Melanesian settlers of New Guinea were infected with Wb after migrating through SE Asia. Then, later the Wb population in PNG was admixed with an ancestral population of Wb infecting the migrating Austronesians 5 kya and eventually infecting Africa. This hypothesis is also supported by human genetic data of admixture between Austronesians and the people of New Guinea, although estimates of proportion vary [Bellwood et al., 2006, Gray et al., 2009, Xu et al., 2012, Skoglund et al., 2016]. The admixture analysis also supports a Wb admixture with the Austronesians in that higher number of clusters (K=4) assign individuals to multiple ancestral populations within either Africa or PNG (supplementary figure S6). The Austronesians or later sea-faring peoples from the Southeast Asia regions [Skoglund et al., 2016] may have carried the same ancestral population of Wb to the Polynesian archipelago. A more formal test of this hypothesis would require comparison of genome sequences of Wb from the Polynesian archipelago with those of PNG and those of Africa (or better Madagascar).

### Austronesian migration spreads *W. bancrofti* to Africa

The Austronesians probably never made it to continental Africa, and are only thought to have colonized Madagascar and the Comoros Islands [Bellwood et al., 2006, Gray et al., 2009]. If the Austronesians carried Wb, then Wb must have first invaded Madagascar. In our study, we did not have samples from Madagascar so can only correlate the divergence time between African populations with that of other migration events. The timing of divergence between African populations of Wb (1.4 –2.5 kya) does coincide with admixture among the Malagasy and Bantu speaking people of Africa (1.8 kya) [Pierron et al., 2014, Pierron et al., 2017] and it is possible that this admixture event spread Wb to continental Africa and subsequently throughout all of Africa.

Finally, we estimated that the ancestral effective population size of the African populations of Wb was 600–1000. This would be expected if the founding population of Wb was one that infected a smaller migrating population. Overall, these results suggest that introduction of Wb into Africa occurred from an initially small population and that Wb then spread and expanded throughout Africa likely from Malagasy or East Africa. Samples from Madagascar and Central African will be required to better resolve this the timing and direction of spread.

### Slave Trade spread *W. bancrofti* to the New World

Our analyses indicate a divergence time between the Haitian and African Wb populations of 0.4-0.6 kya. This timing is consistent with historical records documenting the large number of slaves transported during the transatlantic slave trade from Central and Western Africa (estimates of 4,000 to Jamaica in 1518 AD, [Wynter, 1984]). In contrast to the founding Wb population arriving in Africa, the estimated ancestral effective population size between Haiti and Mali is 8,000 – 32,000, indicating i) that the Wb population had significantly expanded since the initial African colonization and ii) that many more Wb were brought to the New World (possibly through continuous migrations over an extended time).

While the primary admixture results indicate common ancestry between Wb from Haiti with Mali, more subtle analyses (supplementary figure S7) assigned a fraction of the ancestry of Haitian Wb worms to Kenyan Wb populations, possibly owing to multiple Wb admixtures into Haiti. In support of multiple introductions, we find that the Haitian population of Wb has a larger variance in nucleotide diversity than either the Malian or Kenyan Wb populations. Future studies will need to include Wb populations from Central Africa and South America to better identify the sources of admixture.

### Locally adapted haplotypes may affect future drug effectiveness

We identified signals of adaptive selection in separate populations at genes associated with human immune suppression and pesticide sensitivity. Immune suppression may have been critical to avoid endangering hosts in new environments, as high mortality coupled with mal-adapted vectors, would reduce overall parasite survival. Pesticide resistance may be associated with recent pesticide treatments to reduce mosquito populations during malaria control programs or agricultural pests. Since Wb is transmitted by mosquitoes, chemical toxicity may also affect Wb survival. For example, Acetylcholinesterase (AChE) inhibitors are a primary action of carbamates and organophosphates pesticides against arthropods [Casida and Durkin, 2013, Verma et al., 2017]. Selection at genes identified as resistant to AChE inhibitors (first identified in *C. elegans* by [Nguyen et al., 1995]), could, for example, be due to exposure in the vector stage of the Wb life-cycle. Resistance to AChE inhibitors may protect Wb from pesticides used to kill mosquitoes. Future use of AChE inhibitors as an anti-helminthic, while promising in *Trichuris muris* [Sundaraneedi et al., 2018], may have limited efficacy to Wb populations of Mali and Kenya. Tracking changes in allele frequency at these genes during pesticide application may provide further insights on the role of these genes, which would link Wb prevalence to vector control through a novel mechanism of insecticide as a nematicide, but may also endanger future drug development.

Four of the outlier genes have also been identified as potential targets for new anti-parasite drugs in the Malian and Kenyan populations of Wb: Na/Cl GABA transporter [Jiang et al., 2005, Mullen et al., 2006, Casida and Durkin, 2013, O’Neill et al., 2016], PAN domain protein [Kumar et al., 2007, O’Neill et al., 2016], SLO-1 Ca-activated K channel [Welz et al., 2011], phosphatidate cytidylyltransferase [Narayan et al., 1989, Crowther et al., 2010] (table 2). All of our samples were collected prior to mass drug administration; thus, the signal of local adaptation may be in response to other environmental factors rather than widespread drug resistance. Thus, drug resistance could exist in a Wb population before drug treatment has even begun.

## Conclusions

The World Health Organization classifies lymphatic filariasis as a vulnerable infectious disease meaning that it could potentially be completely eradicated. *W. bancrofti* (Wb) is the main causative agent of lymphatic filariasis, so to eliminate lymphatic filariasis we must drive Wb extinct. A better understanding of the genetic diversity and its organization among endemic areas would facilitate tracking recently introduced parasites, identification of possible drug resistance alleles/genes, and relating parasite genomic differences with varying responses to drugs treatments. Our study provides novel tools and data that could help with the agenda. First, our estimates of diversity and relatedness in four endemic Wb populations prior to mass drug administration provide a baseline to evaluate the effectiveness of treatment in each region as well as to identify regions that might need specific renewed efforts. Second, our characterization of genetic polymorphisms provides a list of informative genetic markers that can now be used to efficiently monitor changes in genetic diversity by sequencing amplicons or genotyping (see similar approaches in Plasmodium [Friedrich et al., 2016, Redmond et al., 2017]). Finally, while our data emphasize the limited gene flow between continental populations, they also showed the ability of Wb to successfully migrate between continents and adapt to new environments and vectors and could serve as a cautionary note until all populations are successfully eliminated.

## Methods

### Sample Collection

Samples used in this study from PNG were collected under IRB approved by Case Western Reserve University and PNG – Institute for Medical Research. Haitian samples were collected under IRB protocols reviewed and approved by the CDC IRB and the ethics committee of Hopital Ste. Croix in Leogane, Haiti. The samples from Mali were obtained as part of an NIAID- and the University of Bamko-approved protocol (#02 -I-N200). The samples from Kenya were obtained from the South Kenyan Coast (Msambweni District) as part of a protocol approved by the Kenya Medical Research Institute National Ethical Review Committee and the Institutional Review Board for Human Studies at Case Western Reserve University. Information pertaining to samples and worms are available in supplementary table S2.

### Improvement of *W. bancrofti* genome assembly and annotations

The continuity of the current Wb genome assembly was improved through the addition of PacBio reads. Resulting PacBio sequences were error corrected using proovread v2.06 [Hackl et al., 2014] then used to scaffold the current Wb genome using SSPACE-LongRead [Boetzer and Pirovano, 2014]. The final assembly was gap closed using PBJelly [English et al., 2012]. The improved Wb genome was further scaffolded using Ragout v1.2 [Kolmogorov et al., 2014] with alignments to *B. malayi* (PRJNA10729) and *Loa loa* (PRJNA60051). Sequenced RNA was assembled into transcripts and Maker3 was used to finish annotations. VCFs were annotated using SnpEff v3.4 [Cingolani et al., 2012]. See Supplementary Materials for details on template preparation and sequencing.

### sWGA and Population Sequencing

The program sWGA v0.3.0 [Clarke et al., 2017] was used to design primers for whole-genome amplification of Wb while avoiding amplification of both Human sequences (Hg19) and Wb mitochondrial DNA [Ramesh et al., 2012]. Ten microfilariae (MF) were isolated from each individual blood sample for a total of 200 MF and DNA was isolated following methods in [Small et al., 2016]. After amplification, 42 samples were selected for library preparation and sequencing to include a minimum of 10 samples per geographic location. Primers are available in supplementary table S5, further details on amplification are available in Supplementary Materials.

### Variant calling

Reads passing quality controls were mapped to the improved Wb genome sequence (see data availability) using BWA v0.7.13 [Li and Durbin, 2009]. Variants were called for each individual sample separately using GATK v3.1 HaplotypeCaller v3.5 [McKenna et al., 2010]. The final set of SNPs containing all individuals was filtered to remove putatively repetitive and paralogous sequences (Supplementary Materials).

### Population Structure

Population structure was analyzed using parametric and non-parametric methods: Principal Components Analysis (PCA as implemented in scikit-allel), ADMIXTURE v1.3.0 [Alexander et al., 2009], and phylogeny of genome-wide SNPs [Lee et al., 2014]. The VCF file for structure analysis was pruned to remove SNPs in linkage disequilibrium using modules in the scikit-allel package and an r^2^ threshold of 0.20. Further filters were used to remove singletons, multi-allelic sites, and sites with > 20% missing data resulting in 10,340 positions distributed throughout all the autosomal scaffolds.

ADMIXTURE was run for K (number of ancestral populations) from 2 to 11 with 5-fold cross-validation. Each ADMIXTURE analysis was repeated 10 times with different seeds, resulting in a total of 100 runs for each value of K. In order to better understand the different solutions reported by ADMIXTURE, each value of K was input to the online version of CLUMPAK [Kopelman et al., 2015]. ADMIXTURE was run a second time on the PNG population alone as well as the African and Haitian populations to examine population structure within each cluster.

The program fineSTRUCTURE v2.1.1 [Lawson et al., 2012] was used to explore genetic relatedness among Wb worms for all populations. A VCF file was prepared by phasing SNPs using read-informative phasing available in SHAPEIT2 v2.r837 [Delaneau et al., 2013]. SHAPEIT2 was run in “assemble” mode with the follow options:”–states 200 –window 0.5 –rho 0.000075 –effective-size 14000”. Results were plotted using programs available in the fineSTRUCTURE software package.

### Diversity

Nucleotide diversity and Tajima’s D [Tajima, 1989] were calculated in 10,000 bp non-overlapping windows and plotted for each population using the python package scikit-allel v1.1.0 and Matplotlib v2.0.2. The scaled site-frequency spectrum (multiplied by the scaling factor k * (n – k) / n, where k is the minor allele count and n is the number of chromosomes) was calculated for each population on the LD thinned and folded variant set in scikit-allel. The decay of LD was plotted as the average correlation between genotypes in bins of 100 bp windows using scikit-allel and custom python scripts (https://github.com/stsmall).

### Demography

Estimates of effective population size were calculated using the MSMC2 algorithm [Malaspinas et al., 2016] on individual genomes with 10 bootstrap replicates. MSMC2 input files were constructed using scripts available at msmc-tools (https://github.com/stschiff/msmc-tools), a positive mappability mask (http://lh3lh3.users.sourceforge.net/snpable.shtml) and a negative mask removing sites covered by less than 10 reads in each individual genome. A summary for each population was plotted using the R package ggplot2 after linear extrapolation to synchronize time epochs.

A modified version of PopSizeABC [Boitard et al., 2016], https://github.com/stsmall) was used to estimate the piece-wise changes in effective population size for each Wb population. 500,000 simulation datasets were generated using msprime v0.4.0 with accompanied summary statistics calculated using scripts in PopSizeABC. Residuals and data informativeness were evaluated using the ‘plot’ function in the R package ‘abc’ package v2.1 ([Csilléry et al., 2012]; neural-net option and tolerance of 0.001). Results from PopSizeABC and MSMC2 were combined into a single plot by overlapping co-estimated times using linear extrapolation.

We used *∂*a*∂*i v1.6.3 [Gutenkunst et al., 2009] to model the shared demographic history between pairs of populations. The folded-site-frequency spectrum was used to compute the likelihood of the data under three different models: i) an isolation-with-migration model with symmetrical migration, ii) an isolation-migration model with asymmetrical migration, and iii) a pure isolation model (migration rates set to 0). Each model allowed for the estimate of the ancestral population size as well as the daughter population sizes (modeled as exponential population size change after divergence). Our optimization procedure utilized scripts available from https://github.com/kern-lab/miscDadiScripts.

The block-site-frequency-spectrum was used to compute the likelihoods of multiple population models using the program ABLE v0.1.2 [Beeravolu et al., 2016]. Models assumed stepwise changes in population size under the following scenarios: i) populations diverge without migration, ii) populations diverge followed by asymmetrical migration, iii) populations diverge followed by admixture in the PNG population (supplementary figure S8). Each model was run for using the global search CRS with 50,000 local trees searches. Confidence intervals of parameter estimates were inferred using the methods outlined in [Beeravolu et al., 2016]. Estimates were transformed into year using a mutation rate of 2.9E-9 per generation [Denver et al., 2009] and assuming one generation per year (estimated 8-14 months in [Paily et al., 2009] and in [Farrar et al., 2013]).

### Natural Selection and Local Adaptation

HapFLK was run for each chromosome on the African and Haitian populations with PNG as out-group. Outliers for hapFLK values were calculated using python scripts available at https://forge-dga.jouy.inra.fr/projects/hapflk. Outliers were evaluated using an FDR of 10% in the R package qvalue v2.12.0. We identified the population with the fixed or near fixed allele using local population trees and eigenvalue analysis [Fariello et al., 2013].

Outlier regions were queried against the nr_db database downloaded from NCBI (date of download Jan 31, 2018). Protein sequences were obtained from improved annotations (see above section Reference Genome Improvement). The top five hits, sorted by e-value, were retained and gene ontology inferred by comparison to *B. malayi* [Ghedin et al., 2007] and *C. elegans* genomes.

## Acknowledgements

The authors would like to acknowledge Patrick J. Lammie for contributing samples from Haiti as well as providing comments on the manuscript. This work was supported by grants from the National Institutes of Health (AI103263 to PAZ) and the Clinical and Translational Science Collaborative of Cleveland (UL1TR000439 to PAZ).

## Data Availability

Data is currently being curated and submitted to NCBI and SRA (BAM files, RNA-Seq reads, PacBio reads). This manuscript will be updated with accession numbers as they become available. Until that time the improved *W. bancrofti* assembly, gene annotations, and filtered VCFs are available from the corresponding author upon request.

## Supplementary Methods

### Sample Collection

Haiti samples were provided as cryopreserved peripheral blood mononuclear cells (PBMCs). Mali and Kenya samples were provided as cryopreserved whole blood and later filtered using a 2Nm filter paper to reduce human cells [Erickson et al., 2013]. PNG samples were provided as filtered microfilaria preserved in RNAlater^®^ (Life Technologies) and stored at -20°C. Further sample information can be found in Supp Table S1.

### Improvement of the Wb draft genome sequence and gene annotation

Pac Bio sequencing using chemistry RS II was performed on DNA isolated from the sample(s) described in [Small et al., 2016]. Following whole genome amplification (as noted in [Small et al., 2016]), the whole genome amplified DNA was treated according to methods in [Zhang et al., 2006] to reduce chimeric errors. DNA was cleaned using a 1.6:1 Agencourt AMPure XP bead concentration and then eluted in 20ul ddH2O water. DNA was then incubated in a T100(TM) Thermocycler (Bio-Rad) with 1ul of Phi29 enzyme with 25uM concentration of dNTPs and 1x BSA for 30 minutes at 30 °C followed by 65 °C for 3 minutes to deactivate the Phi29 enzyme. DNA was once again cleaned with 1.6:1 Agencourt AMPure XP bead concentration and incubated with one unit of S1 Nuclease to cleave junctions of branched DNA molecules. DNA was again cleaned (see Above) and nick-repaired using PreCR(R) DNA Repair (New England Biolabs, Inc.) and then quantified by Qubit Flourometric Quantitation (ThermoFisher Scientific). The DNA template was prepared and sequenced by the McGill University and Génome Québec. Ten PacBio SMRT cells were run for an estimated coverage of 20-40X. Whole genomes were aligned using progressiveCactus (https://github.com/glennhickey/progressiveCactus) with a guide tree reconstructed from whole mitochondrial genomes [Small et al., 2014]. Ragout [Kolmogorov et al., 2014] was run allowing for repeat resolution with scaffolds named according to *B. malayi* reference genome.

RNA was isolated from a single PNG sample preserved in RNALater. Template and library preparation followed the standard TruSeq RNA Kit TM protocol (Illumina, Inc.). The RNA library was sequenced at 100 base pairs as paired-end reads at the Case Western Reserve University Genomics Core on an Illumina HiSeq 2500. Resulting reads were mapped to the Human genome reference 19 (Hg19) as well as the Wb genome (PRJNA275548) using HISAT [Kim et al., 2015] to separate reads belonging to the human host and then Wb worm. Sequences mapping successfully to Wb and not human were used in Maker3 [Cantarel et al., 2008] along with Wb EST libraries (SAW95SjL-WbMf) and protein sequences curated for *B. malayi* (PRJNA10729) and *Loa loa* (PRJNA60051). Maker3 was run for three progressive iterations using initial gene prediction based on single copy orthologs identifies using the program BUSCO[Simão et al., 2015].

### sWGA and Population Sequencing

Primer sets were tested by amplifying DNA isolated from one infected-patient blood sample as well as from two single microfilaria (MF). DNA quality and ploidy were confirmed by amplifying and sequencing the mitochondrial cytochrome oxidase I (CO1) gene [Small et al., 2013]. Primer sets were nearly indistinguishable using metrics of read depth and percent of the genome covered [Clarke et al., 2017, Leichty and Brisson, 2014], so the larger primer set (12 primers, supplementary table S5) was selected to obtain more even coverage for fragmented DNA.

Isolated DNA was used directly for whole genome amplification following the protocol in [Leichty and Brisson, 2014], except with 1.5 *µ*g of each of 12 custom primers. The reaction was incubated for 8 hours at 30 °C followed by a 70 °C for 15min in T100(TM) Thermocycler (Bio-Rad). Reaction time was chosen to maximize DNA yield while minimizing excessive duplication after analyzing reactions of 4, 6, 8, 10, and 12 hours. After 8-hour incubation, amplified samples were cleaned using a 1.6:1 Agencourt AMPure XP bead concentration to remove small fragments. 40 cycles of quantitative PCR (qPCR) using SYBR^®^ (Life Technologies) in a Mastercycler^®^RealPlex2 (Eppendorf) were used to determine the relative proportions of host (human) and parasite (Wb) DNA. Human DNA was quantified using a custom designed primer pair to amplify a section of Chromosome I:HuQPCR1-F 5’-ACTTTGGGTCATTCCCACTG-3’, HuQPCR1-R 5’ -GCTCAGCTCCTTGCTGGATA -3’. Wb DNA was quantified using primers to amplify isotype-1 of the *β*-tubulin gene [Hoti et al., 2003]. Overall success rate of amplification, measured by final DNA concentration (> 500 ng), from single microfilaria (MF) was 60%.

A 1.5-2 *µ*g of amplified DNA was used in the TruSeq DNA PCR-Free kit (Illumina, Inc.) following steps for the 550 bp insert size. Finished libraries were quantified on a 2100 Bioanalyzer Instrument (Agilent). Samples were pooled in equal molarity and then initially sequenced on Illumina MiSeq for 35 bp to rigorously quantify the proportion of Wb and human DNAs in each library as well as library complexity before final sequencing. Resulting sequencing were mapped to the Wb genome using BWA [Li and Durbin, 2009] and used to adjust pooling ratios between samples. A total of 12 pooled libraries (each containing 6 sample libraries) were sequenced on 12 lanes of the Illumina HiSeq X for 150 base pairs in paired-end mode at the McGill University and Génome Québec. Sequencing resulted in a total of 1.2 Tb of sequencing data from 42 sample libraries.

### Quality control

All sequences were first trimmed for adapter sequences using TrimGalore (http://www.bioinformatics.babraham.ac.uk/ and then examined for base-quality using fastqc (https://www.bioinformatics.babraham.ac.uk/projects/fastqc/) and multiqc (https://github.com/ewels/MultiQC). Filtered read data were then mapped to the mitochondrial genome of Wb [Ramesh et al., 2012] to verify that each sample contained DNA from only a single genome. Three samples were discarded for excess heterozygosity found on the mitochondrial genome (supplementary table S2) and four samples for excess missing data.

### Variant Calling

GATK v3.1 HaplotypeCaller [McKenna et al., 2010] was run with the following parameters: “-newQual –emitRefConfidence GVCF –pcr_indel_model NONE”. The resulting gVCF from HaplotypeCaller were converted to VCF format using GATK program GenotypeGVCFs. VCF positions were removed if they failed the following filter criteria: “QD < 5, QUAL < 30, DP < 14, MQ < 30, MQRankSum < -12.5, ReadPosRankSum < -8.0, FS > 60.0, ABHet < .30, ABHet > 0.70, ABHom < 0.90”. Positions passing all filters in each individual sample were merged to produce a final set of SNPs across all samples. As combining SNPs among samples can lead to missing data, homozygous or uncalled in original sample, the candidate SNP set was then used in freebayes to fill homozygous positions [Garrison and Marth, 2012]. Putatively repetitive and paralogous sequences were identified using RepeatMasker [Tarailo-Graovac and Chen, 2009] and Snpable (http://lh3lh3.users.sourceforge.net/snpable.shtml).

## Supplementary Tables and Figures

**Figure S1:**
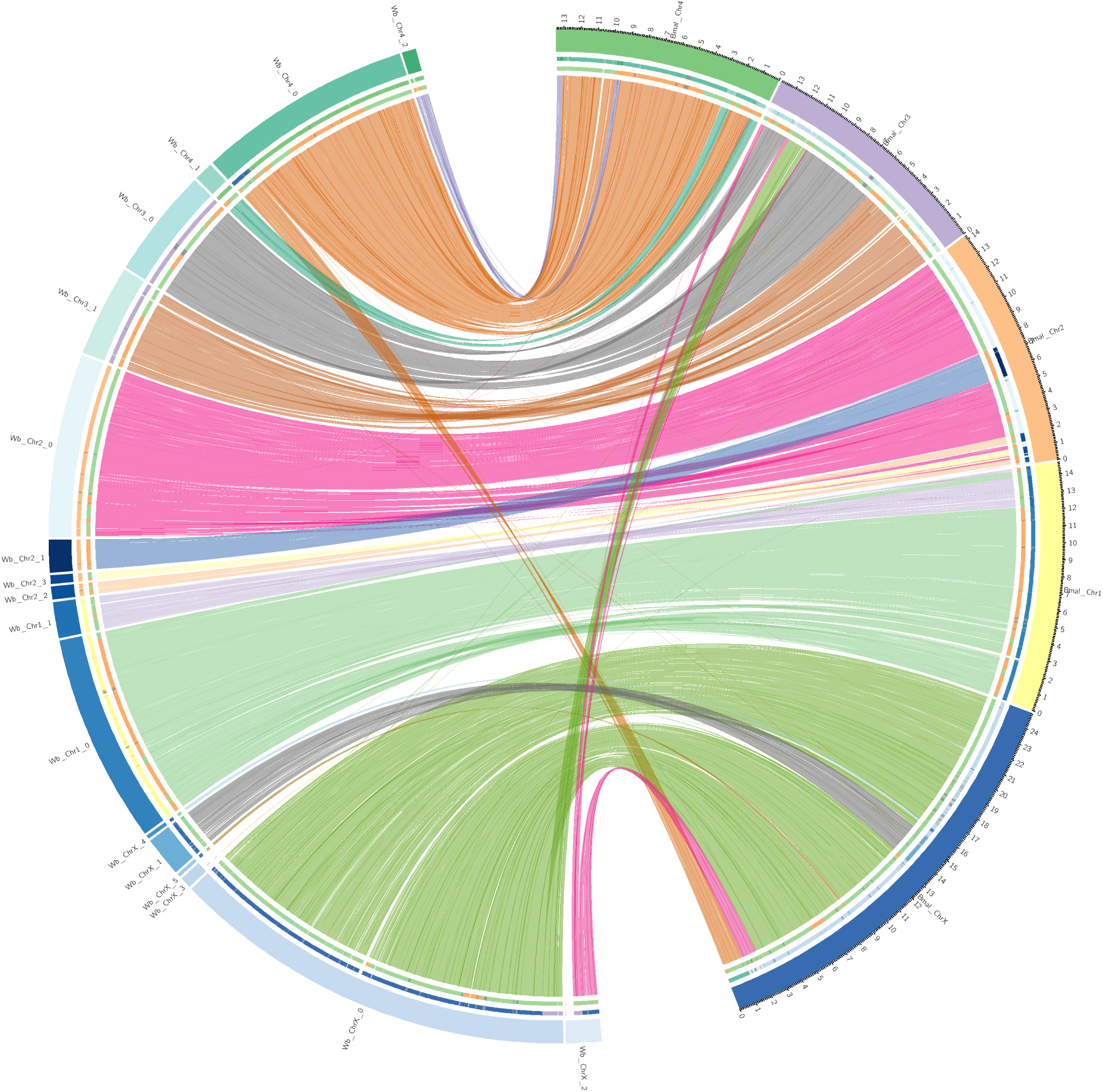
Circos plot comparing *W. bancrofti* with *B. malayi.*

**Figure S2:**
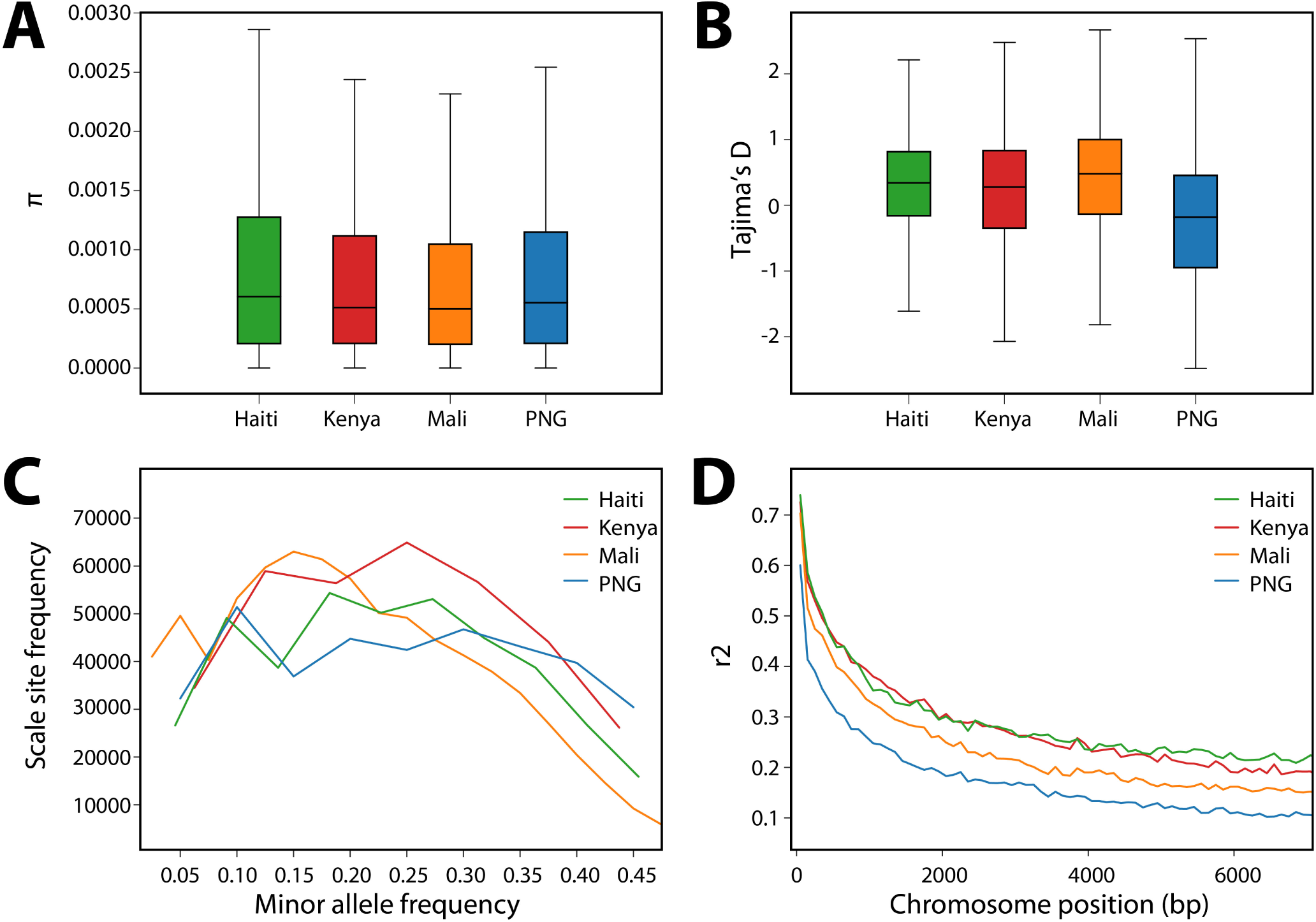
Diversity.

**Figure S3:**
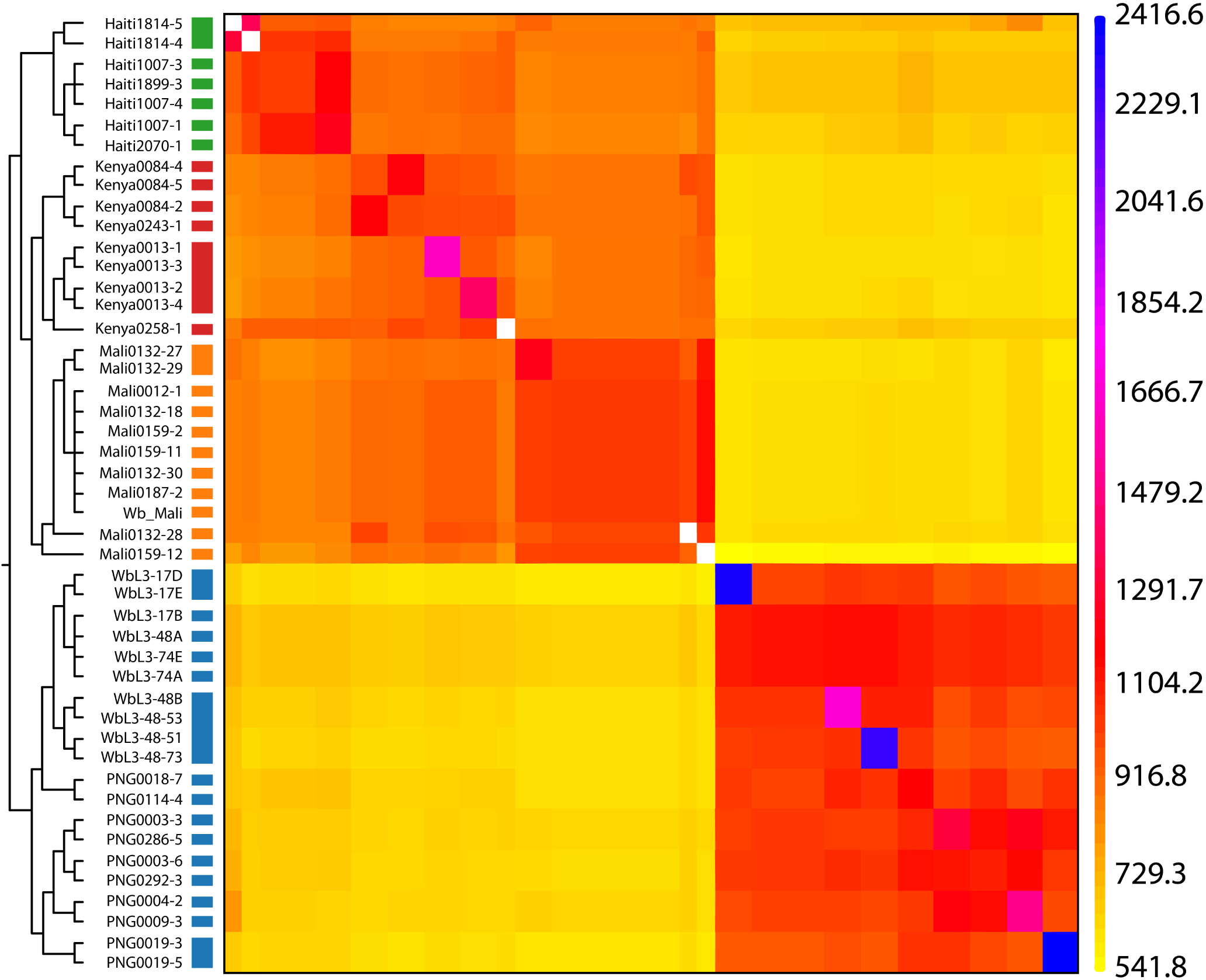
fineSTRUCTURE.

**Figure S4:**
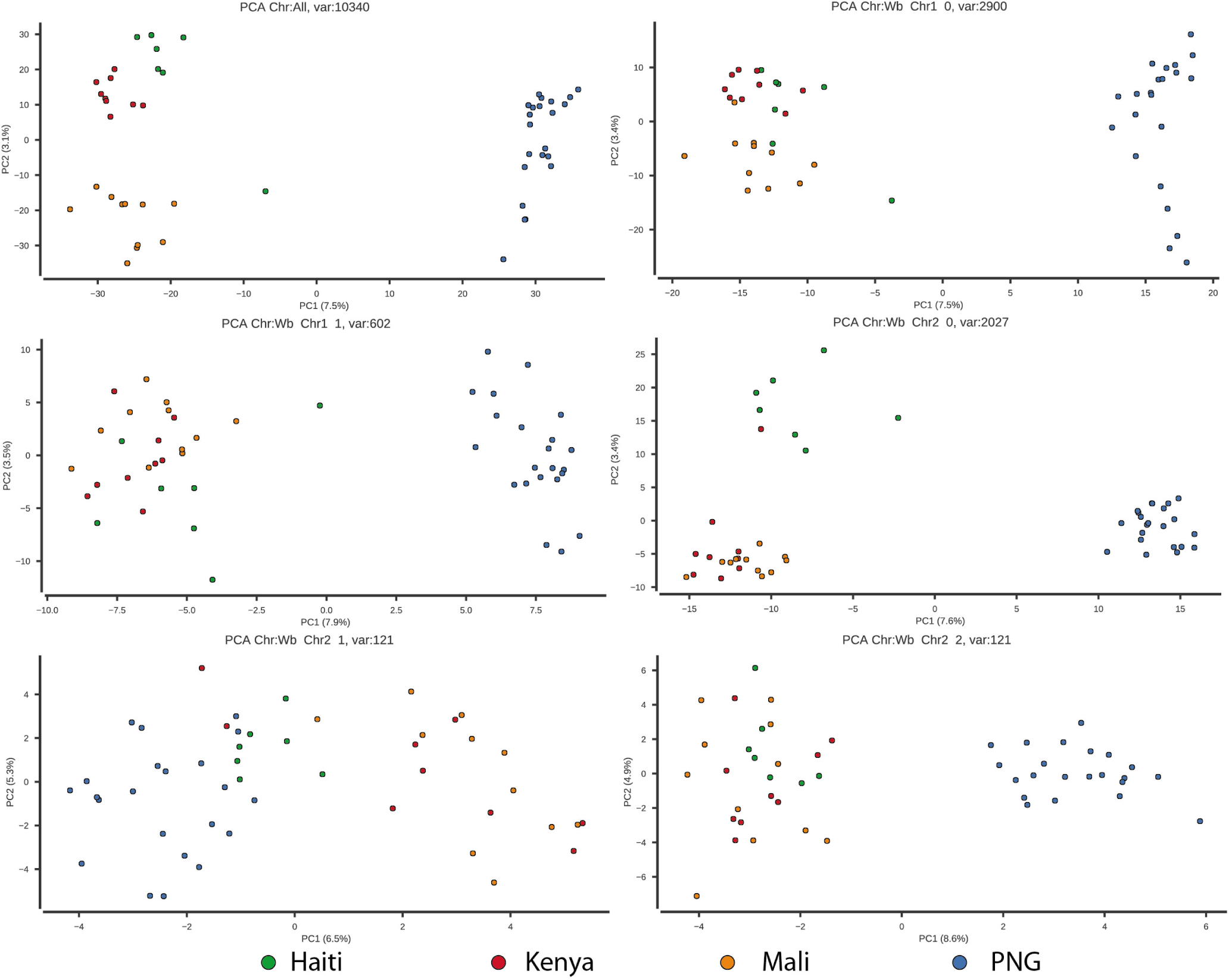
PCA Chromosomes.

**Figure S5:**
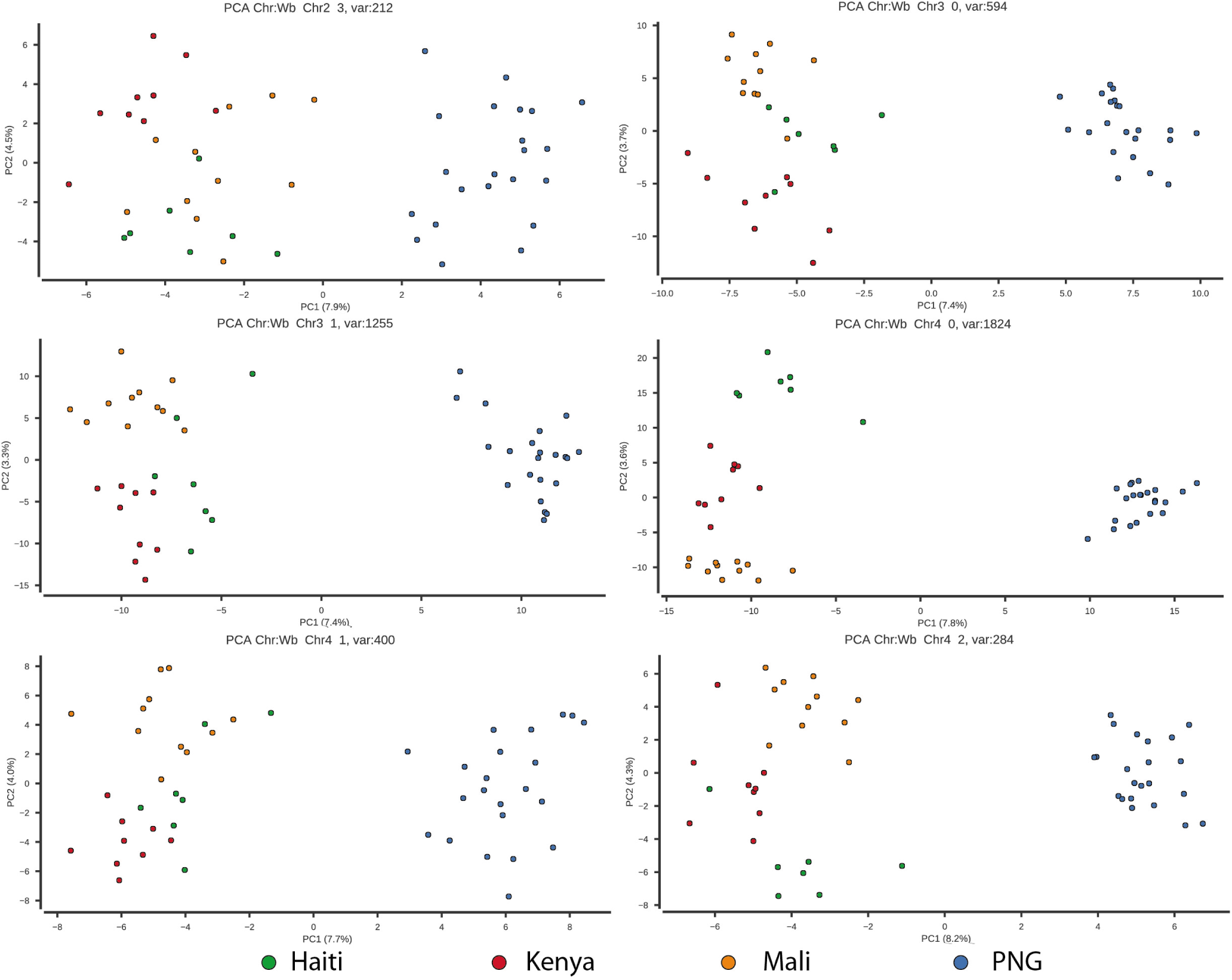
PCA Chromosomes.

**Figure S6:**
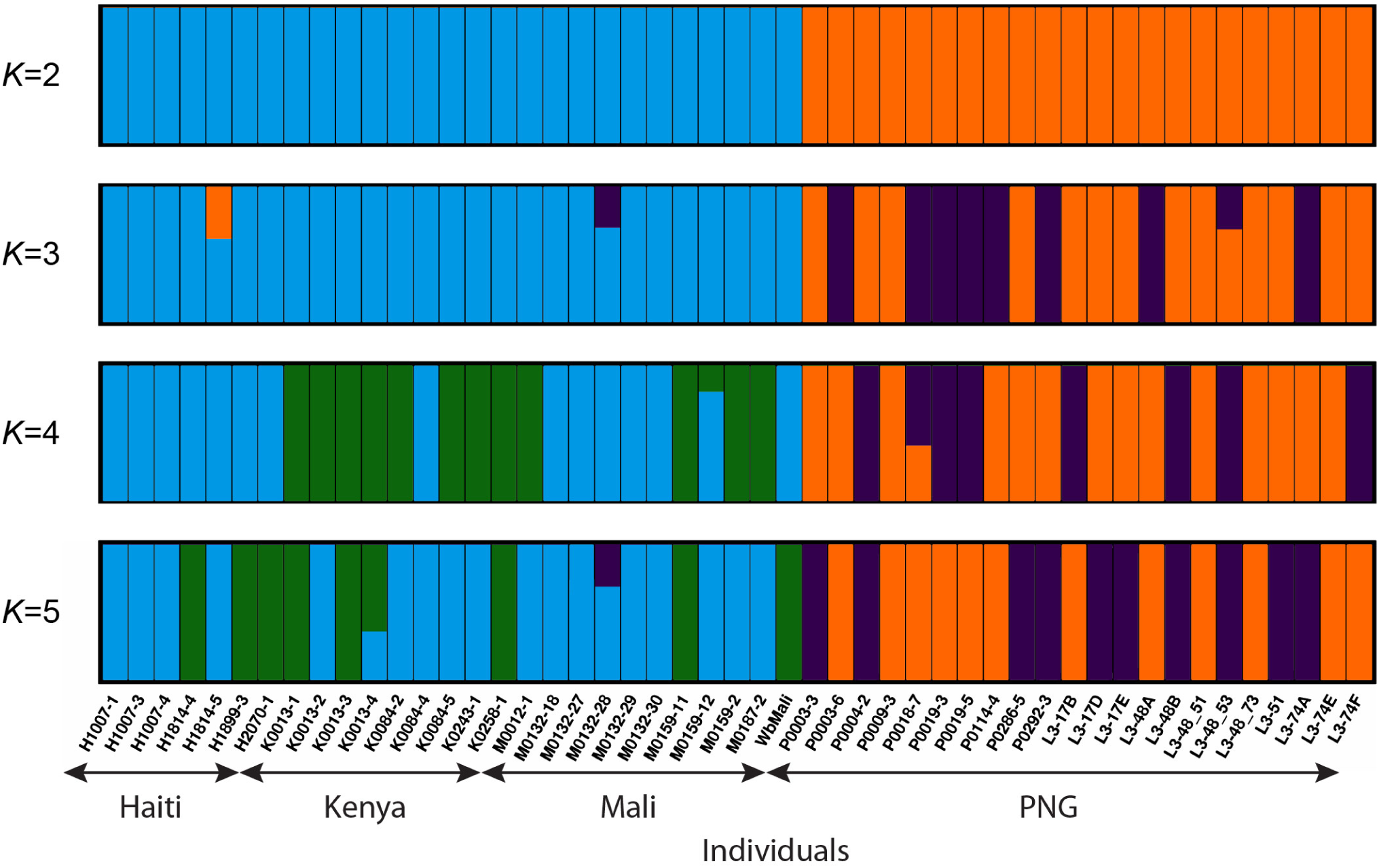
Admixture.

**Figure S7:**
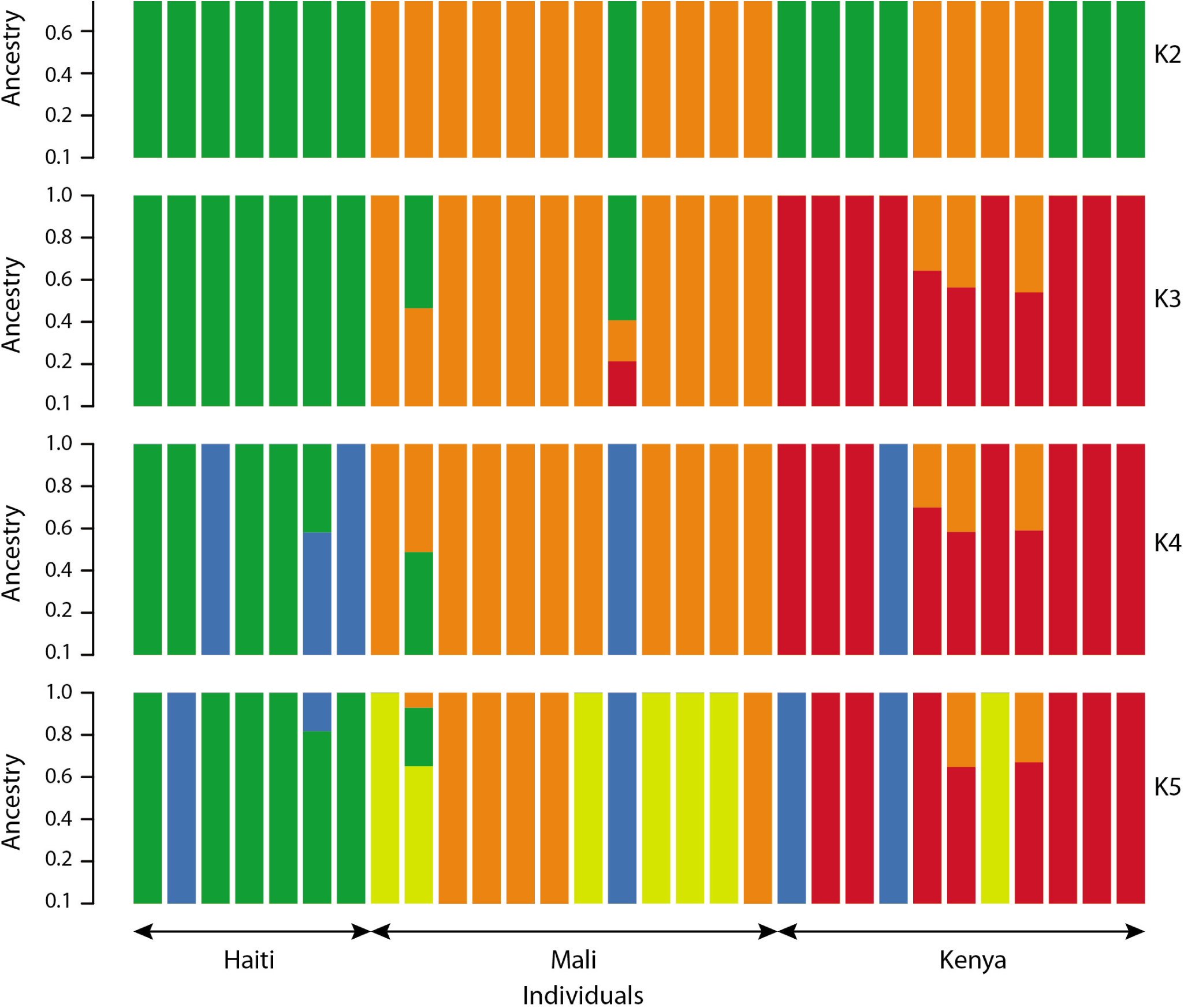
Admixture Haiti-Mali-Kenya.

**Figure S8:**
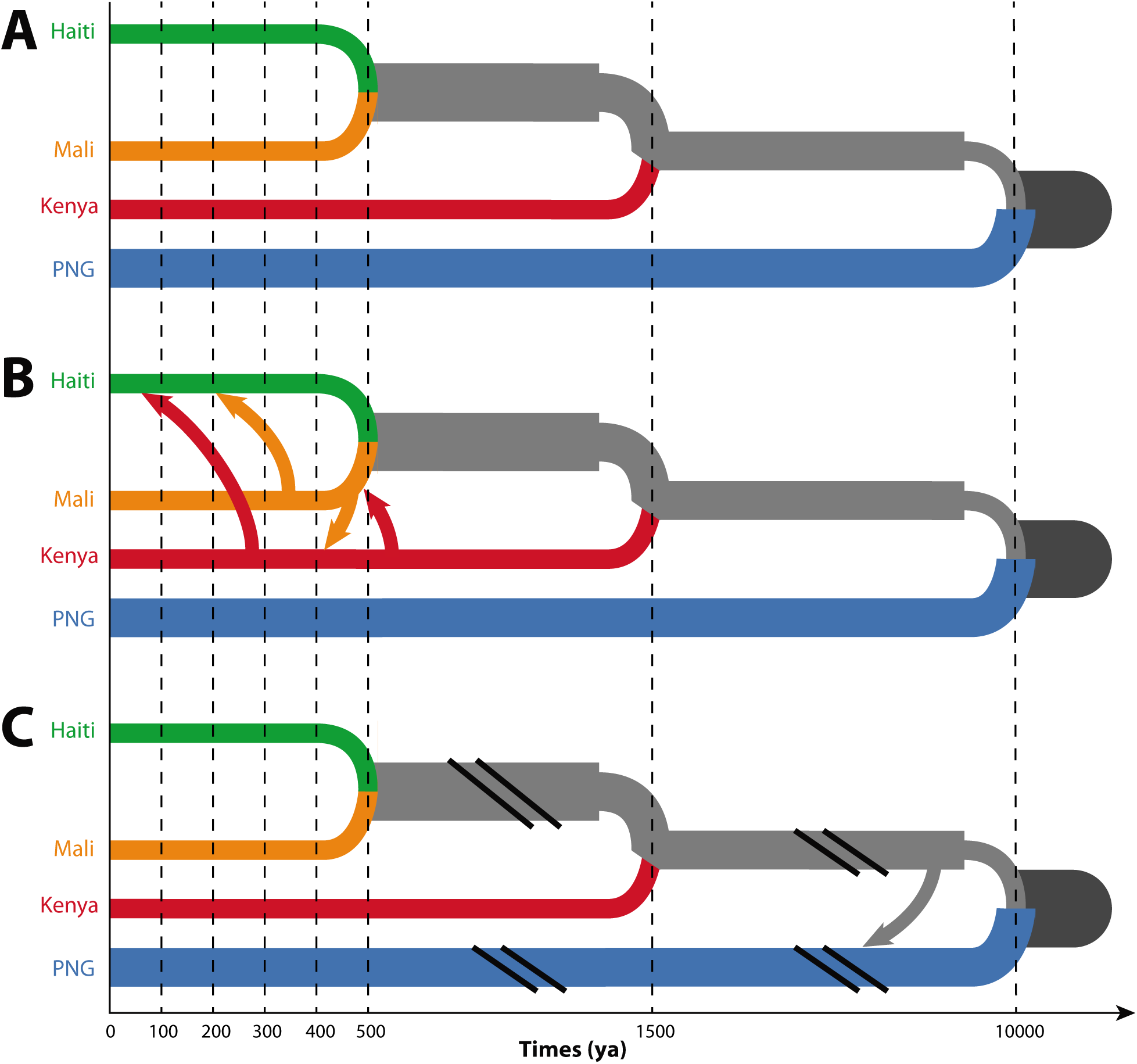
Admixture Haiti-Mali-Kenya.

**Table S1:** Assembly Results.

| Chromosome | Scaffold ID | Size | Ns | Genes | Gene density |
| --- | --- | --- | --- | --- | --- |
| 1 | Wb_Chr1_0 | 12,368,652 | 81,951 | 1,633 | 0.000132 |
|  | Wb_Chr1_1 | 2,119,897 | 98,244 | 182 | 0.000086 |
| 2 | Wb_Chr2_0 | 10,841,590 | 87,666 | 1,143 | 0.000105 |
|  | Wb_Chr2_1 | 1,918,629 | 415 | 253 | 0.000132 |
|  | Wb_Chr2_2 | 750,293 | 0 | 106 | 0.000141 |
|  | Wb_Chr2_3 | 490,737 | 2,531 | 58 | 0.000118 |
| 3 | Wb_Chr3_0 | 6,490,679 | 17,498 | 917 | 0.000141 |
|  | Wb_Chr3_1 | 5,401,749 | 67,144 | 569 | 0.000105 |
| 4 | Wb_Chr4_0 | 12,704,880 | 207,068 | 1,369 | 0.000108 |
|  | Wb_Chr4_1 | 1,128,067 | 610 | 98 | 0.000087 |
|  | Wb_Chr4_2 | 813,737 | 109,849 | 45 | 0.000055 |
| X | Wb_ChrX_0 | 24,139,319 | 571,287 | 2,710 | 0.000112 |
|  | Wb_ChrX_1 | 2,355,808 | 45,023 | 210 | 0.000089 |
|  | Wb_ChrX_2 | 2,085,878 | 19,869 | 137 | 0.000066 |
|  | Wb_ChrX_3 | 662,921 | 28,893 | 66 | 0.000100 |
|  | Wb_ChrX_4 | 204,088 | 0 | 5 | 0.000024 |
|  | Wb_ChrX_5 | 173,113 | 0 | 16 | 0.000092 |

**Table S2:**
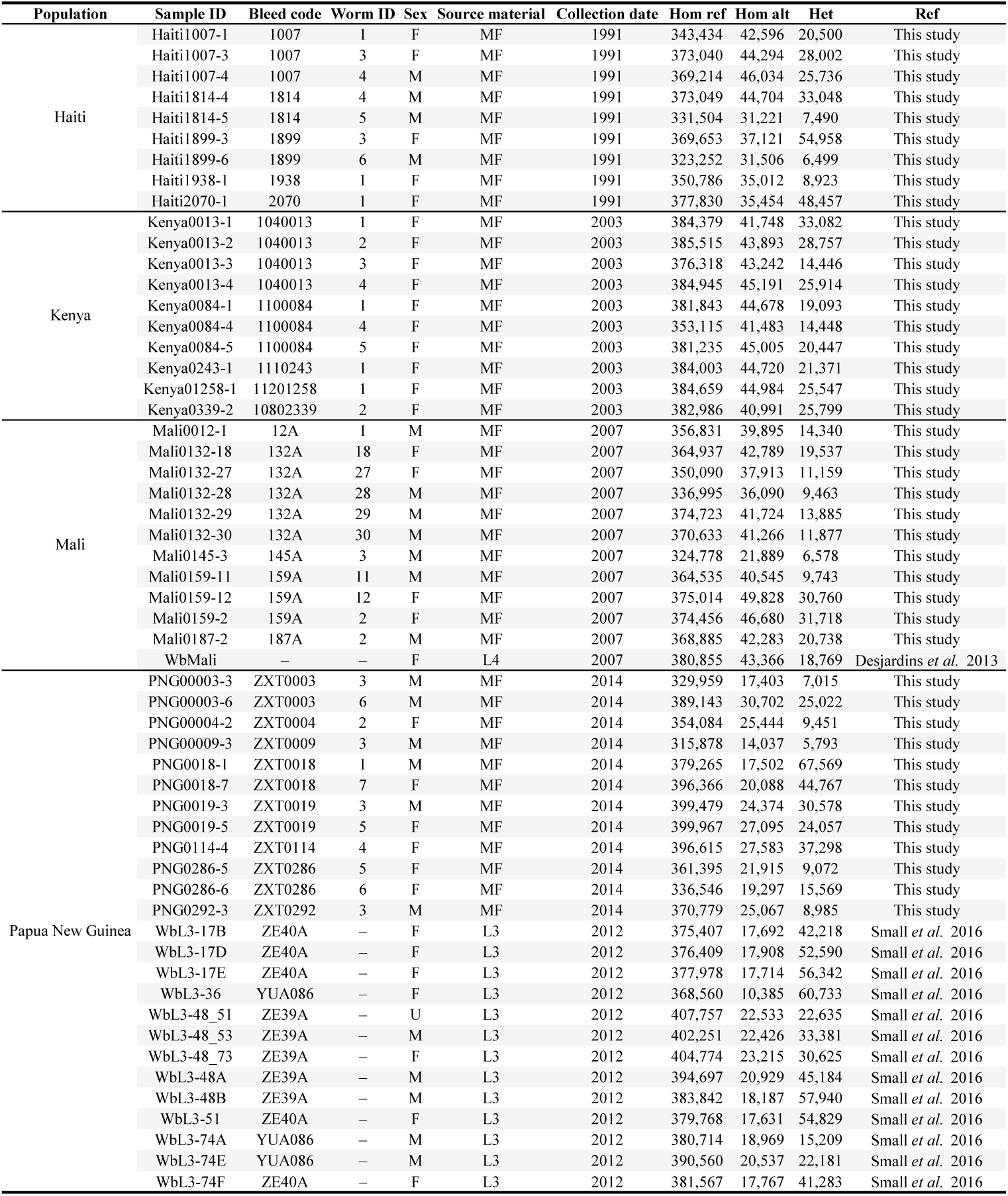
Samples.

**Table S3:** Variants.

| Population | SNPs | Singletons | Doubletons | Private |
| --- | --- | --- | --- | --- |
| Haiti | 165412 | 43168 | 33663 | 24776 |
| Mali | 152458 | 31910 | 27479 | 23201 |
| Kenya | 149685 | 33863 | 28585 | 23564 |
| PNG | 204138 | 51289 | 31812 | 78091 |

**Table S4:**
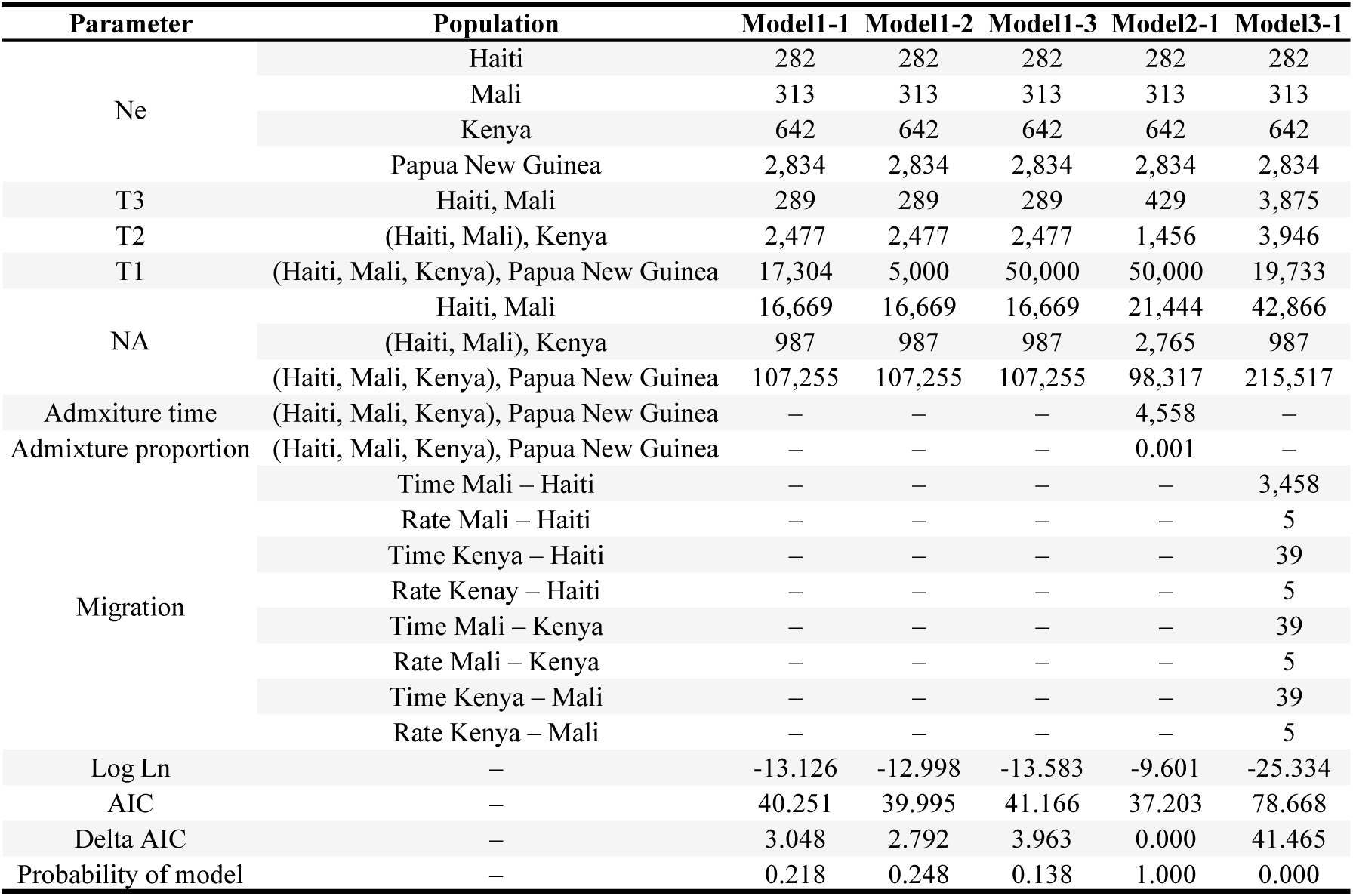
Model Results.

| Parameter | Population | Model1-1 | Model1-2 | Model1-3 | Model2-1 | Model3-1 |
| --- | --- | --- | --- | --- | --- | --- |
| Ne | Haiti | 282 | 282 | 282 | 282 | 282 |
|  | Mali | 313 | 313 | 313 | 313 | 313 |
|  | Kenya | 642 | 642 | 642 | 642 | 642 |
|  | Papua New Guinea | 2,834 | 2,834 | 2,834 | 2,834 | 2,834 |
| T3 | Haiti, Mali | 289 | 289 | 289 | 429 | 3,875 |
| T2 | (Haiti, Mali), Kenya | 2,477 | 2,477 | 2,477 | 1,456 | 3,946 |
| T1 | (Haiti, Mali, Kenya), Papua New Guinea | 17,304 | 5,000 | 50,000 | 50,000 | 19,733 |
| NA | Haiti, Mali | 16,669 | 16,669 | 16,669 | 21,444 | 42,866 |
|  | (Haiti, Mali), Kenya | 987 | 987 | 987 | 2,765 | 987 |
|  | (Haiti, Mali, Kenya), Papua New Guinea | 107,255 | 107,255 | 107,255 | 98,317 | 215,517 |
| Admixture time | (Haiti, Mali, Kenya), Papua New Guinea | — | — | — | 4,558 | — |
| Admixture proportion | (Haiti, Mali, Kenya), Papua New Guinea | — | — | — | 0.001 | — |
| Migration | Time Mali – Haiti | — | — | — | — | 3,458 |
|  | Rate Mali – Haiti | — | — | — | — | 5 |
|  | Time Kenya – Haiti | — | — | — | — | 39 |
|  | Rate Kenya – Haiti | — | — | — | — | 5 |
|  | Time Mali – Kenya | — | — | — | — | 39 |
|  | Rate Mali – Kenya | — | — | — | — | 5 |
|  | Time Kenya – Mali | — | — | — | — | 39 |
|  | Rate Kenya – Mali | — | — | — | — | 5 |
| Log Ln | — | -13.126 | -12.998 | -13.583 | -9.601 | -25.334 |
| AIC | — | 40.251 | 39.995 | 41.166 | 37.203 | 78.668 |
| Delta AIC | — | 3.048 | 2.792 | 3.963 | 0.000 | 41.465 |
| Probability of model | — | 0.218 | 0.248 | 0.138 | 1.000 | 0.000 |

**Table S5:** sWGA Primers.

| Primer | Sequence |
| --- | --- |
| swga01 | AATCGATA*A*T |
| swga02 | ACGAATAA*T*T |
| swga03 | CGACGA*A*T |
| swga04 | CGATAC*G*A |
| swga05 | CGCGAA*A*A |
| swga06 | CGTAAA*C*G |
| swga07 | GACGAAAA*A*A |
| swga08 | TCGAAC*G*A |
| swga09 | TCGCGA*A*A |

